# Knowledge-guided deep learning models of drug toxicity improve interpretation

**DOI:** 10.1101/2022.02.28.482300

**Authors:** Yun Hao, Joseph D. Romano, Jason H. Moore

**Affiliations:** Genomics and Computational Biology (GCB) Graduate Program, University of Pennsylvania, Philadelphia, PA; Institute for Biomedical Informatics, University of Pennsylvania, Philadelphia, PA; Department of Computational Biomedicine, Cedars-Sinai Medical Center, Los Angeles, CA

## Abstract

In drug development, a major reason for attrition is the lack of understanding of cellular mechanisms governing drug toxicity. The black-box nature of conventional classification models has limited their utility in identifying toxicity pathways. Here we developed DTox (**D**eep learning for **Tox**icology), an interpretation framework for knowledge-guided neural networks, which can predict compound response to toxicity assays and infer toxicity pathways of individual compounds. We demonstrate that DTox can achieve the same level of predictive performance as conventional models with a significant improvement in interpretability. Using DTox, we were able to rediscover mechanisms of transcription activation by three nuclear receptors, recapitulate cellular activities induced by aromatase inhibitors and PXR agonists, and differentiate distinctive mechanisms leading to HepG2 cytotoxicity. Virtual screening by DTox revealed that compounds with predicted cytotoxicity are at higher risk for clinical hepatic phenotypes. In summary, DTox provides a framework for deciphering cellular mechanisms of toxicity *in silico*.

## INTRODUCTION

With the application of quantitative high-throughput screening techniques, toxicity testing programs^1,2^ have generated millions of data points regarding the response of biological systems to important chemical libraries, both *in vitro* and *in vivo*. Specifically, in the Tox21 program^1^, over 8,500 compounds were tested for a variety of toxicity endpoints including stress response, genotoxicity, cytotoxicity, developmental toxicity, etc. These toxicity profiles can assist with probing how chemicals interact with proteins and pathways to trigger a certain outcome, and thus shed light on cellular mechanisms of toxicity^3^. Furthermore, with the help of machine learning algorithms, researchers can identify the chemical or biological patterns of a compound that might be predictive of adverse health outcomes in human^4,5^.

Previous studies have modeled toxicity endpoints from physiochemical properties of compound using a wide range of supervised learning algorithms, including *k*-nearest neighbors^6,7^, Bayesian matrix factorization^8^, support vector machines^7,9^, random forests^6,10^, gradient boosting^11^, and more recently, deep neural networks^12–14^. Even though most of these algorithms achieved decent predictive performance, none of them could overcome the trade-off between accuracy and interpretability. As algorithmic design gets more complex, it becomes challenging to interrogate how each input feature contributes to the eventual prediction^15^. A few post-hoc explanation techniques, such as local interpretable model-agnostic explanations (LIME)^16^ and deep learning important features (DeepLIFT)^17^, were developed to address the challenge. Nevertheless, these techniques often draw criticism in that they only provide an approximate explanation with locally fitted naïve models. Thus, they may not reflect the real behavior of original model^18^. More critically, the setting of existing toxicity prediction models has limited the explanation of contributions from structural properties or target proteins while interactions with pathways remain largely uncharacterized. For toxicologists, the behavior of pathways proves crucial in deciphering the cellular activities induced by a compound, and understanding how target proteins, specific pathways, and biological processes trigger the toxicity outcome as a whole^5^. Therefore, a toxicity prediction model that achieves interpretability at both the gene and pathway level is urgently needed.

Recent developments in visible neural networks (VNN) have overcome the accuracy-interpretability trade-off. VNN is a type of neural network whose structure is guided by extensive knowledge from biological ontologies and pathways. The incorporation of ontological hierarchy in VNN forms a meaningful network structure that connects input gene features to output response via hidden pathway modules, making the model highly interpretable at both gene and pathway level. In a pioneering study, Ma *et al*. built a VNN with 2,526 Gene Ontology and Clique-eXtracted Ontology terms, for predicting growth rate of yeast cells from gene deletion genotypes^19^. The authors were also able to rediscover key ontology terms responsible for cell growth by examining the structure of the VNN. Subsequent studies have extended the VNN model for learning tasks regarding human cells, such as predicting drug response and synergy in cancer cell lines^20^, modeling cancer dependencies^21^, and stratifying prostate cancer patients by treatment-resistance state^22^. It is our working hypothesis that VNN can address the limitations of existing toxicity prediction models due to its incorporation of pathway knowledge and the resulting high interpretability. In this study, we employed the Reactome^23^ pathway hierarchy to develop a VNN model—namely DTox—for predicting compound response to 15 toxicity assays. Further, we developed a DTox interpretation framework for identifying VNN paths that can explain the toxicity outcome of compounds. We connected the identified VNN paths to cellular mechanisms of toxicity by showing their involvement in the target pathway of respective assay, their differential expression in the matched LINCS experiment^24^, and their compliance with screening results from mechanism of action assays. We applied the DTox models of cell viability to perform a virtual screening of ~700,000 compounds and linked the predicted cytotoxicity score with clinical phenotypes of drug-induced liver injury. We concluded with a discussion of potential discoveries made by DTox, some of which have already been validated in previous studies. Our code and data can be accessed openly at https://github.com/yhao-compbio/DTox. In general, the DTox interpretation framework will benefit *in silico* mechanistic studies and generate testable hypotheses for further investigation.

## RESULTS

### Design and training of DTox VNN models for predicting compound response to toxicity assays

We designed a VNN structure (**Fig. 1; Methods**) that connects target proteins (input features) to assay outcome (output response) via Reactome pathways (hidden modules). The feature profile containing 361 target proteins was inferred from structural properties of a compound using our previously developed method, TargetTox^25^. By our design, each pathway is represented by 1-20 neurons depending on its size. Connections between input features and the first hidden layer are constrained to follow protein-pathway annotations while the connections among hidden layers are constrained to follow child-parent pathway relations. The incorporation of pathway hierarchy makes DTox VNN model highly interpretable, in contrast to conventional black-box neural network models.

**Figure 1.**
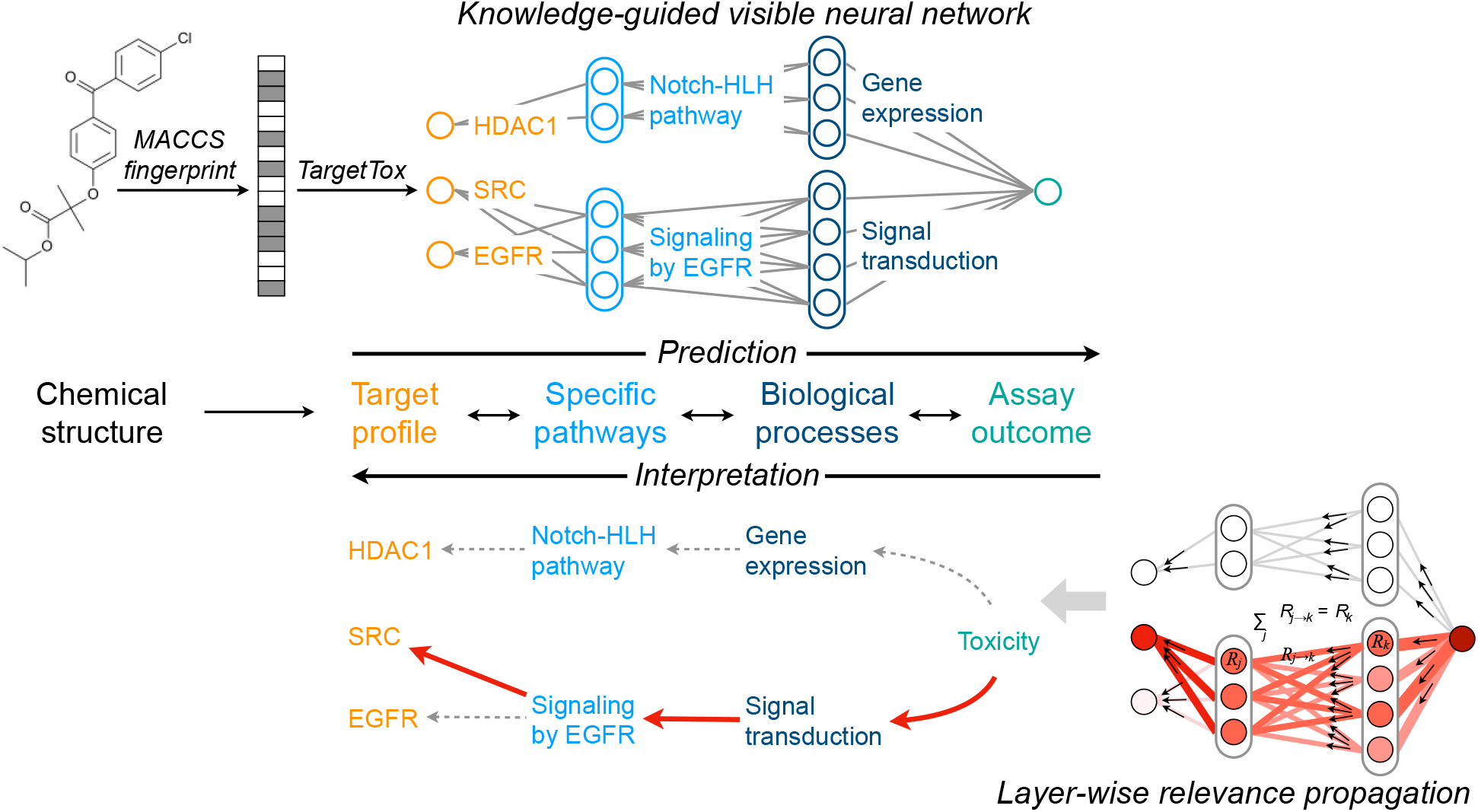
Modeling compound response to toxicity assay with DTox. For toxicity prediction, the chemical structure of a compound is quantified using MACCS fingerprints before being converted to target profile by our previously developed method, TargetTox. The target profile is then fed into our DTox VNN, whose structure is guided by Reactome pathway hierarchy. Specific pathways and biological processes are coded as hidden modules with a series of neurons. For model interpretation, the network output is propagated backward onto each neuron as relevance score using the layer-wise relevance propagation technique. A permutation-based strategy is then employed to identify the VNN paths of high relevance. Each path connects a compound to its toxicity outcome via the target protein, specific pathways, and biological process.

We trained DTox VNN models on 15 datasets from the Tox21 high throughput screening program^1^. Each dataset contains active and inactive compounds for one toxicity assay, with an average of 5,178 available compounds per assay (**Supplementary Fig. 1a; Methods**). We implemented an early stopping criterion to speed up the training process (**Methods**). As a result, the training process was completed within 100-200 epochs for all 15 datasets (**Supplementary Fig. 2**). We also implemented hyperparameter tuning by grid search to derive an optimal model for the prediction of each assay outcome (**Supplementary Table 1**; **Methods**). On average, an optimal DTox VNN model contains 412 hidden pathway modules (**Supplementary Fig. 1b**), and 45,623 neural network parameters (**Supplementary Fig. 1c**). The average ratio between number of training samples versus number of network parameters is 0.13±0.03 (**Supplementary Fig. 1d**), with the model of estrogen receptor agonist assay being the highest (0.31) and the model of hedgehog antagonist assay being the lowest (0.07). Compared to a conventional multi-layer perceptron (MLP) model, DTox VNN model has far fewer network parameters. On average, the number of network parameters for a DTox VNN only accounts for three percent of the number for a matched MLP (**Supplementary Fig. 1e**).

To customize the network structure for prediction of each assay outcome, we made the root biological process a hyperparameter of DTox VNN (**Methods**). This means through hyperparameter tuning, we can choose a branch or combination of branches from the Reactome pathway hierarchy that result in the best predictive performance for an assay of interest (**Fig. 2a**). For instance, signal transduction pathways alone can deliver the optimal model for HEK293 cell viability assay while additional pathways from immune system are required for HepG2 cell viability prediction, suggesting a potential role of immune response in HepG2 cytotoxicity. In general, models built with multiple branches perform better than models built with a single branch.

**Figure 2.**
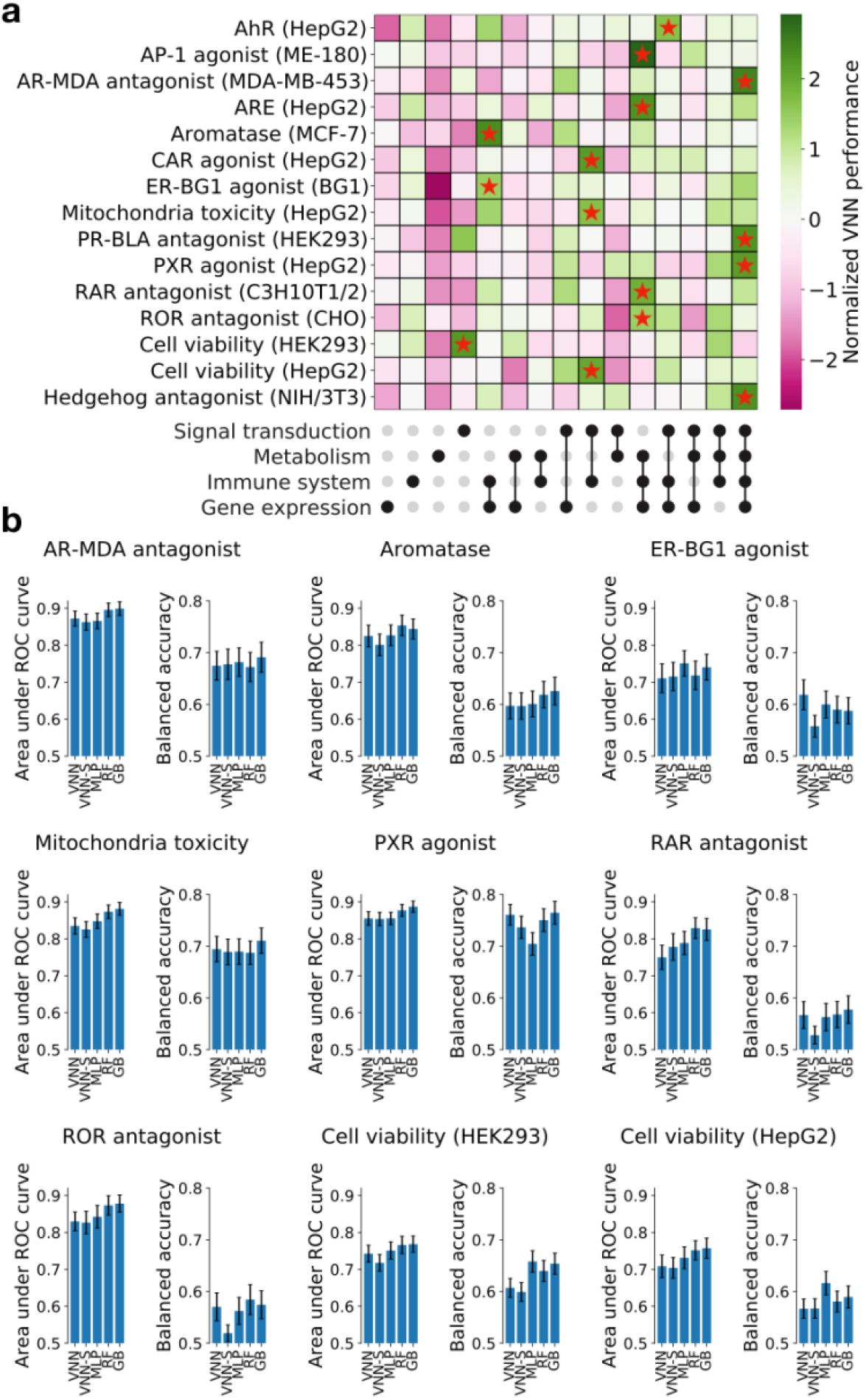
Prediction of compound response to 15 toxicity assays. (**a**) Heatmap showing the training performance of DTox VNN built under different combinations of root biological processes (shown as upset plot at the bottom). To facilitate comparison, the model performance is normalized within each assay using Z-transform. The optimal combination for each assay is highlighted with a red star. The name of each assay is annotated on the left, with name of the assay cell line included in parenthesis. AhR: aryl hydrocarbon receptor, AP-1: activator protein-1, AR-MDA: androgen receptor in MDA-kb2 AR-luc cell line, ARE: antioxidant response element, CAR: constitutive androstane receptor, ER-BG1: estrogen receptor in BG1 cell line, PR-BLA: progesterone receptor in PR-UAS-bla HEK293T cell line, PXR: pregnane X receptor, RAR: retinoid acid receptor, ROR: retinoid-related orphan receptor. (**b**) Barplot showing the validation performance in nine toxicity assays (see **Supplementary Figure 3** for the remaining six assays). The nine selected assays are involved in the subsequent analyses. Performance of DTox VNN (VNN) is compared against four other models: an alternative visible neural network built with shuffled pathway hierarchy (VNN-S), a multi-layer perceptron with the same number of hidden layers and neurons as DTox VNN (MLP), random forest (RF), and gradient boosting (GB). Performance is measured by two metrics: area under ROC curve and balanced accuracy, with error bar shows the 95% confidence interval.

### DTox VNN can achieve the same level of performance as complex classification algorithms

We validated the predictive performance of DTox VNN models on held-out validation sets, which on average contain 1,295 compounds per assay. The optimal models of all 15 assays exhibit an area under the ROC curve (AUROC) greater than 0.7 (0.7-0.8: 6 models, 0.8-0.9: 9 models). Similarly, 14 models exhibit a balanced accuracy above 0.55 (0.55-0.65: 9 models, 0.65-0.75: 4 models, > 0.75: 1 model) except for the optimal model of AP-1 signaling agonist assay. We then compared the optimal performance of DTox VNN against four other classification algorithms (**Fig. 2b** and **Supplementary Fig. 3**; **Supplementary Table 2**; **Methods**). Comparing DTox VNN to an alternative model built with shuffled pathway hierarchy (VNN-S), we observed two assays where DTox VNN significantly outperformed VNN-S in balanced accuracy (estrogen receptor agonist and retinoid-related orphan receptor gamma antagonist). Comparing DTox VNN to a matched MLP model, we observed one assay where DTox VNN significantly outperformed MLP in balanced accuracy (pregnane X receptor agonist), and two assays in the opposite direction (HEK293 and HepG2 cell viability). Comparing DTox VNN to random forest (RF) and gradient boosting (GB), we observed one assay where DTox VNN significantly outperformed both RF and GB (constitutive androstane receptor agonist), and one assay in the opposite direction (AP-1 signaling agonist). In general, DTox VNN model achieved the same level of predictive performance as these well-established classification algorithms.

### Development of a DTox interpretation framework for explaining VNN predictions

A fundamental advantage of VNN over other classification algorithms lies in its high interpretability. The incorporation of pathway hierarchy enables us to reason through hidden layers of VNN for mechanistic interpretation. Therefore, we developed a DTox interpretation framework to identify paths from VNN that can explain the toxicity outcome of a compound (**Fig. 1**; **Methods**). Each identified path links together a root biological process, its descendant pathway modules, and a target protein feature. The framework has two hyperparameters: *γ* and *ε*. *γ* controls the stability of interpretation results while *ε* controls the sparsity. We evaluated the effect of hyperparameter settings on identified VNN paths (**Supplementary Fig. 4**; **Methods**). We observed that the set of identified paths exhibits consistently high similarity across distinct hyperparameter settings, as the average Jaccard Index reaches 0.70. Due to the high similarity, we only used the VNN paths identified from one setting (*γ* = 0.001, *ε* = 0.1) for the following validation analyses (**Supplementary Table 3**).

### DTox interpretation framework can rediscover mechanisms of transcription activation by nuclear receptor

To evaluate whether DTox interpretation framework can rediscover known mechanism for a toxicity outcome, we looked for “ground truth” from the VNN paths identified for four nuclear receptor assays: androgen receptor antagonist, estrogen receptor agonist, retinoic acid receptor antagonist, and retinoid-related orphan receptor gamma antagonist. Each of the four assays measures compound response to a specific nuclear receptor transcription pathway. Therefore, we established ground truth as the VNN path that links together root process of gene expression, nuclear receptor transcription pathway, and the specified target receptor (AR, ERα, RARβ, RORγ; **Fig. 3**). In three of the four nuclear receptor assays, our framework was able to identify the ground truth path for at least 29% of all active compounds (ERα: 29%, RARβ: 31%, RORγ: 54%). Comparing to identifying by chance, our framework improved the proportion by at least two-folds.

**Figure 3.**
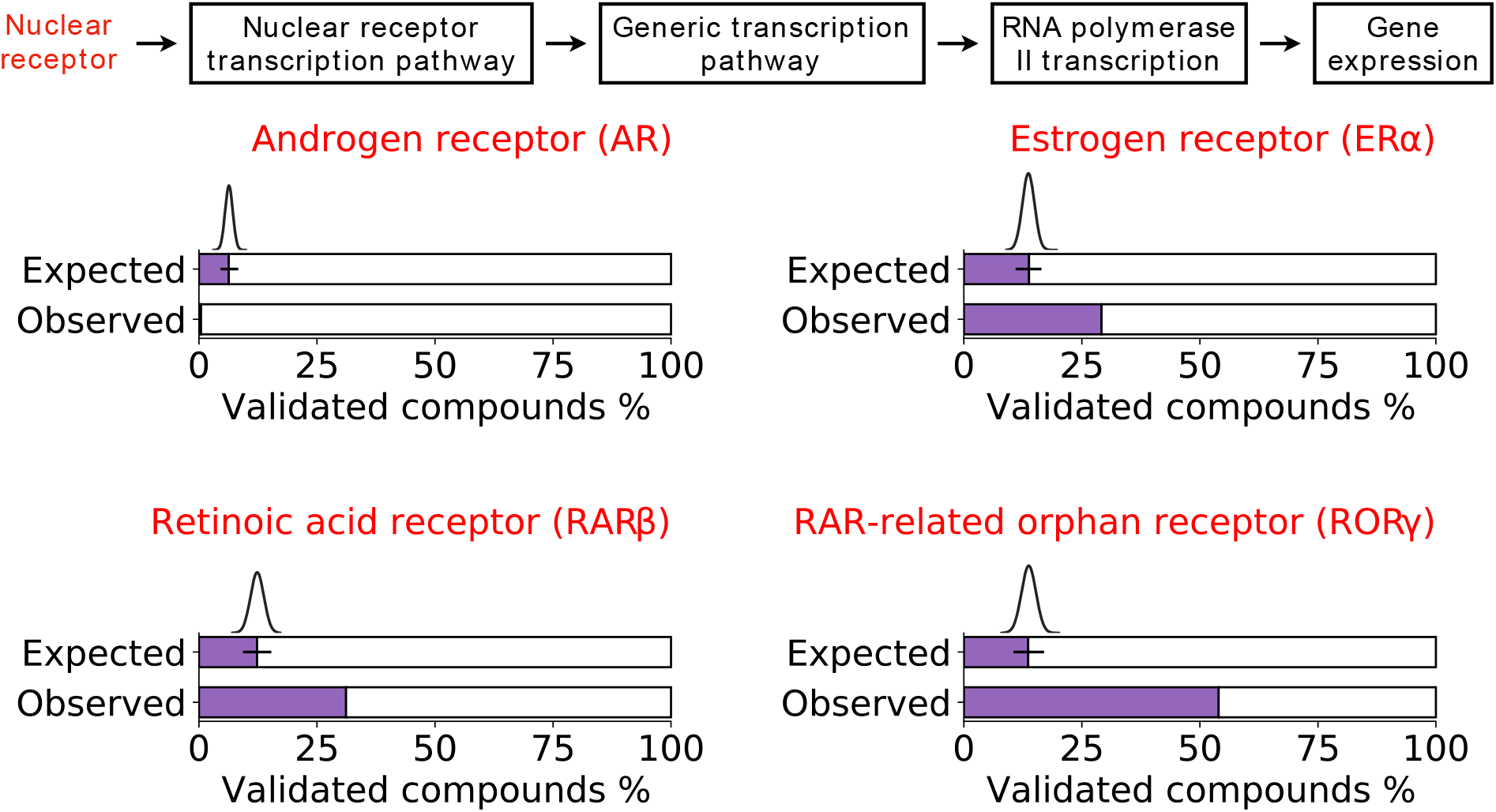
Validation of identified VNN paths by known mechanisms. The “ground truth” VNN path placed at the top represents the known mechanism of transcription activation by nuclear receptor. A compound is considered to be validated by the known mechanism if the ground truth path is identified by DTox. The validation analysis is performed for four nuclear receptor assays, with each barplot comparing the observed versus expected proportion of validated compounds for a single receptor. The expected proportion is computed by random sampling, with the histogram and fitted density curve showing the sampled distribution (95% confidence interval shown as error bar).

### DTox interpretation framework can recapitulate cellular activities induced by aromatase inhibitors and pregnane X receptor agonists

To evaluate whether DTox interpretation framework can recapitulate cellular activities induced by active compounds, we studied the differential expression of VNN paths identified for four assays: aromatase inhibitor, mitochondria toxicity, pregnane X receptor agonist, and HepG2 cell viability (**Methods**). In total, we obtained the gene expression profile measured from 321 LINCS experiments in which an active compound was used to treat the assay cell line (121 experiments for aromatase inhibitor assay, 54 for mitochondria toxicity assay, 101 for pregnane X receptor agonist assay, and 45 for HepG2 cell viability assay). Of all 321 experiments, we found 161 (50%) cases where DTox interpretation framework was able to identify at least one differentially expressed VNN path. On average, 3.8±0.6% of VNN paths identified by our framework were found to be differentially expressed, significantly higher than the expected proportion by chance (2.4±0.3%, *P* = 3.5e^-5^). We then performed the comparison separately by assay and dose-time combinations (**Fig. 4a**). In aromatase inhibitor assay, our framework outperformed the expected proportion across all three dose-time combinations (FDR = 0.01). In pregnane X receptor agonist assay, our framework outperformed the expected proportion among the two groups of experiments conducted 24 hours after treatment (FDR = 0.03). In HepG2 cell viability assay, although no overall difference was detected among experiments conducted 6 hours after treatment (FDR = 0.07), our framework was still able to identify a relatively high proportion of differentially expressed VNN paths for individual compounds such as cilnidipine (21.4%), cyclopamine (12.5%), and chloroxine (10%).

**Figure 4.**
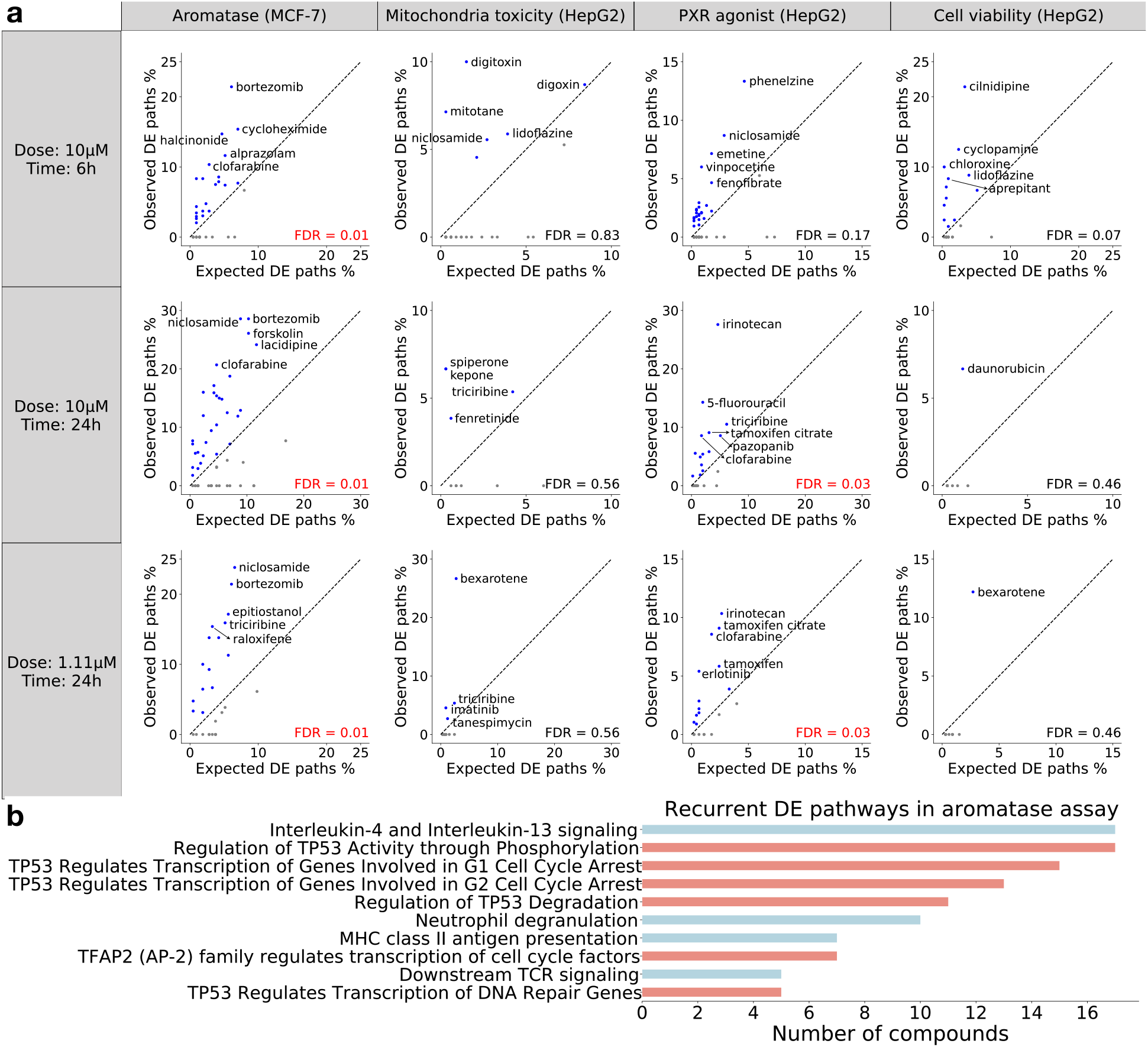
Validation of identified VNN paths by differential expression. (**a**) A VNN path is considered to be differentially expressed (DE) if all pathways along the path is enriched for DE genes from the matched LINCS experiment. The validation analysis is performed for four assays (columns) in three different dose-time groups (rows), with each scatter plot comparing the observed versus expected proportion of DE paths for a single group. The observed proportion is computed with VNN paths identified for each compound while the expected proportion is computed with all possible VNN paths. A T-test is employed to examine whether the average observed proportion of each group is significantly higher than the average expected proportion (FDR value shown at the bottom right). The diagonal is shown as black dashed line, with compounds in the upper triangle (observed > expected) shown in blue and compounds in the lower triangle (observed < expected) shown in grey. Compounds with the top five observed proportion in each group are annotated with their names. (**b**) Barplot showing the DE VNN paths that are recurrently identified for at least five aromatase inhibitors. Each VNN path is named after its lowest-level pathway. Paths that contain the “transcriptional regulation by TP53” pathway is highlighted in salmon while the remaining paths are colored in cyan.

Based on the results of differential expression analysis, induced cellular activities appear to be more consistent among aromatase inhibitors compared to the other three assays. As we discovered ten differentially expressed VNN paths that are recurrently identified for at least five aromatase inhibitors (**Fig. 4b**). By contrast, we only discovered one such VNN path for the other three assays combined. Interestingly, “transcriptional regulation by TP53” and its descendant pathways are involved in six of the ten discovered VNN paths, suggesting a potential mechanism for regulation of aromatase by p53 in the MCF-7 aro estrogen responsive element (MCF-7aro/ERE) breast cancer cell line, a finding supported by previous study^26^. In addition to p53, interlukin-4 and interlukin-13 also appears to play an important role in regulation of aromatase, as the relevant VNN path is linked to 17 aromatase inhibitors by differential expression. The finding is worth further experimental investigation.

### DTox interpretation framework can differentiate distinctive mechanisms leading to HepG2 cytotoxicity

Next, we sought to explain the compound-induced cytotoxicity in HepG2 cells using VNN paths identified for HepG2 cell viability assay. A recent review paper^27^ summarized four major mechanisms leading to cell death in drug-induced liver injury (DILI): (i) TNFR1/2 mediated apoptosis via caspase activation and pro-survival inhibition, (ii) MST1/2 mediated apoptosis via Hippo signaling, (iii) immune response activation via MHC class II antigen presentation, and (iv) TLR3/4 mediated necrosis (**Fig. 5a**). Since HepG2 cell line was derived from liver tissue, we can use the four mechanisms as a reference for compound-induced cytotoxicity in HepG2 cells. We identified nine Reactome pathways that participate in the four mechanisms (**Fig. 5a**). We then mapped HepG2-cytotoxic compounds to the nine Reactome pathways via VNN paths identified by our framework (**Supplementary Fig. 5**). Of all 1,120 cytotoxic compounds, 707 (63%) compounds are mapped to at least one of the nine cell death-related pathways. We performed hierarchical clustering on the mapping and identified two compound clusters (**Supplementary Fig. 5**). Compounds in first cluster are linked to cytotoxicity via apoptosis while compounds in the second cluster are linked to cytotoxicity via immune activation and necrosis. Nevertheless, we discovered a few compounds that exhibit characteristics of both clusters. For instance, according to our framework, mifepristone, a medical abortion drug, causes cytotoxicity in HepG2 cells by activating both apoptosis (via MAPK1/MAPK3 signaling and Hippo signaling) and necrosis (via TLR3 and TLR4 cascade), a finding supported by previous studies^28–30^ (**Fig. 5d**). In addition, our framework was able to link mifepristone with its therapeutic target, glucocorticoid receptor, via PTK6 signaling (**Supplementary Table 3).** The other therapeutic target of mifepristone, progesterone receptor, is not in the VNN.

**Figure 5.**
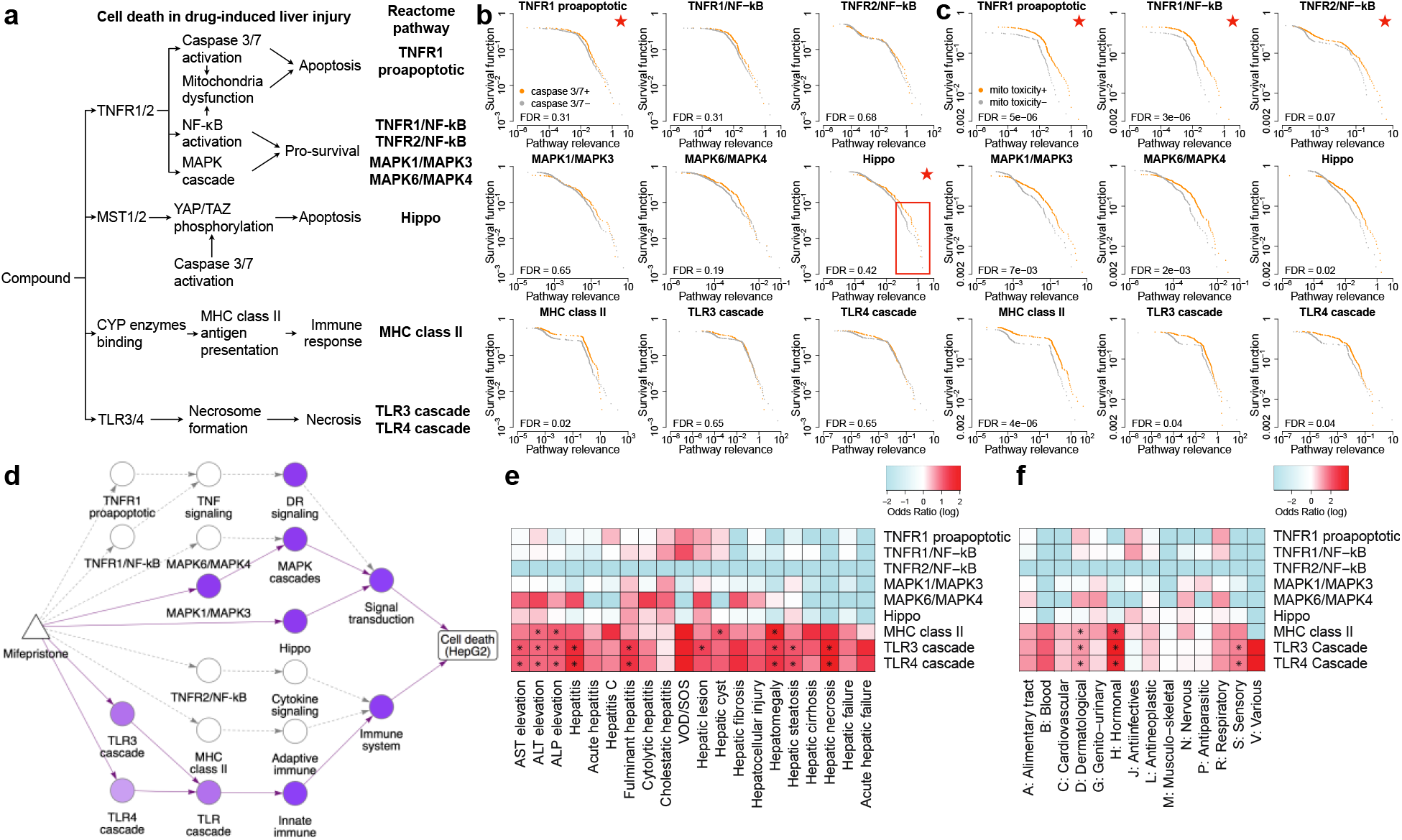
In-depth analysis of HepG2 cytotoxicity using identified VNN paths. (**a**) Established mechanisms for cell death in drug-induced liver injury. Reactome pathways relevant to the mechanisms are identified and used as reference for the analysis. (**b** and **c**) Survival plots comparing the pathway relevance scores among active (orange curve) versus inactive (grey curve) compounds of two mechanisms of action assays: caspase 3/7 induction (**b**) and disruption of the mitochondrial membrane potential (**c**). Comparisons are made for nine cell death-related pathways, with each plot showing the comparison for a single pathway. Red star at the top right denotes that the pathway is related to the respective mechanism of action. Log-rank test is employed to examine whether the two distributions in each plot are significantly different (FDR value shown at the bottom left). (**d**) Network diagram showing the simplified DTox VNN structure connecting mifepristone (triangle node) to the HepG2 cytotoxicity (rectangle node) via pathway modules (round nodes). Pathways with relevance score > 0 are colored in purple, with the scale proportional to relevance scale. The VNN paths identified for mifepristone by DTox are shown in solid lines while the rest are shown in dashed lines. (**e** and **f**) heatmaps showing the enrichment of nine cell death-related pathways among compounds associated with 20 drug-induced liver injury phenotypes (**e**) and among compounds of 14 ATC classes (**f**). Cells are colored based on odds ratio. Fisher’s exact test is employed to examine the significance of enrichment (asterisk denotes FDR < 0.05). VOD/SOS: veno-occlusive disease and sinusoidal obstruction

To evaluate whether DTox interpretation framework can differentiate distinctive mechanisms leading to HepG2 cytotoxicity, we looked for accordance between the assigned pathway relevance and screening results from two mechanism of action assays (included in the Tox21 datasets). The first assay we studied measures caspase 3/7 induction in HepG2 cells. Caspase 3 and caspase 7 are key executioners of apoptosis^31^. They are involved in TNFR1/2 mediated apoptosis, and YAP/TAZ phosphorylation of Hippo signaling^32^ (**Fig. 5a**). Accordingly, we compared the assigned relevance scores between caspase 3/7+ and caspase 3/7-compounds regarding TNFR1-induced proapoptotic signaling and Hippo signaling (**Fig. 5b**). Overall, we did not observe significantly higher relevance among caspase 3/7+ compounds regarding the two signaling pathways (FDR = 0.31 and 0.42, respectively).

However, for Hippo signaling, we did observe higher pathway relevance among caspase 3/7+ compounds above the 90^th^ percentile of two distributions (highlighted in **Fig. 5b**), hence a partial agreement between assigned pathway relevance and caspase 3/7 induction screening. By contrast, the pattern among top-ranked compounds was not observed in other cell death-related pathways except for MHC class II antigen presentation (FDR = 0.02; **Fig. 5b**), suggesting a potential role of caspase 3/7 in MHC class II antigen presentation, a finding worth of further investigation.

The second assay we studied measures disruption of the mitochondrial membrane potential (MMP). MMP is a key indicator of mitochondrial activity as it is required for ATP synthesis. Disruption of MMP can lead to release of cytochrome c, which in turn amplifies the apoptosis signal^33^. The downstream effectors of TNFR1/2, including caspase activation and inhibition of NF-κB activation, can cause disruption of MMP^33,34^ (**Fig. 5a**). Accordingly, we compared the assigned relevance scores between MMP-disrupting and nondisruptive compounds regarding TNFR1-induced proapoptotic signaling, TNFR1-induced NF-κB signaling, and TNFR2-induced NF-κB signaling (**Fig. 5c**). We observed significantly higher relevance among MMP-disrupting compounds regarding TNFR1-induced proapoptotic and TNFR1-induced NF-κB signaling (FDR = 5e^-6^ and 3e^-6^, respectively). And the pattern of higher relevance is consistent across all percentiles of two distributions, hence an agreement between assigned pathway relevance and MMP disruption screening. By contrast, the pattern was not observed in other cell death-related pathways except for MHC class II antigen presentation (FDR = 4e^-6^; **Fig. 5c**), suggesting the potential involvement of mitochondria in antigen presentation, a finding supported by previous study^35^.

### Interpretation of HepG2 cytotoxicity links clinical phenotypes of DILI to TLR3/4 mediated necrosis

We also sought to explain 20 clinical phenotypes of DILI using the derived mapping between compounds and nine cell death-related pathways. For each DILI phenotype, we identified the enriched pathways among its associated compounds (**Fig. 5e**; **Methods**). We observed a disproportionate prevalence of high odds ratio in the two necrosis-related pathways (TLR3 and TLR4 cascade signaling) across almost all DILI phenotypes, with hepatic necrosis, hepatitis, and hepatic fibrosis being the three highest. In total, nine phenotypes are significantly enriched for TLR3/4 mediated necrosis (FDR < 0.05). By contrast, only four phenotypes are significantly enriched for immune activation via MHC class II antigen presentation while no phenotype is significantly enriched for Hippo signaling or TNFR1/2 mediated apoptosis. These results suggest that TLR3/4 mediated necrosis is a common cause for clinical phenotypes of DILI, a finding supported by previous studies^36,37^.

Similarly, we identified the enriched pathways among compounds of 14 Anatomical Therapeutic Chemical (ATC) classes (**Fig. 5f**). Each ATC class represents a group of drugs that act on a specific organ or system. We found three classes (hormonal, sensory, and dermatological) significantly enriched for TLR3/4 mediated necrosis, and two classes (hormonal and dermatological) significantly enriched for immune activation via MHC class II antigen presentation.

### HepG2 cytotoxicity score predicted by DTox VNN can differentiate hepatic cyst compounds from negative controls

Finally, we implemented the optimal DTox VNN models of two cell viability assays (HepG2 and HEK293) to predict the probability of cytotoxicity for 708,409 compounds from DSSTox^38^ (**Supplementary Table 4**). The list of compounds provides a great coverage of the chemical landscape of interest to toxicological and environmental researchers. To demonstrate the clinical application of DTox VNN, we sought to differentiate DSSTox compounds associated with DILI phenotypes from negative controls using the predicted HepG2 cytotoxicity score (**Fig. 6**). We were able to detect significantly higher predicted scores among the compounds associated with hepatic cyst (*P* = 0.015), as hepatic cyst is the only DILI phenotype showing a significant association with HepG2 cytotoxicity (*OR* = 1.90, 95% CI: 1.04-3.45). Among the remaining 19 DILI phenotypes showing weak or no association with HepG2 cytotoxicity (9 phenotypes with OR > 1, 10 with OR < 1), we were only able to detect a significant difference for one phenotype: hepatic steatosis (*P* = 0.008).

**Figure 6.**
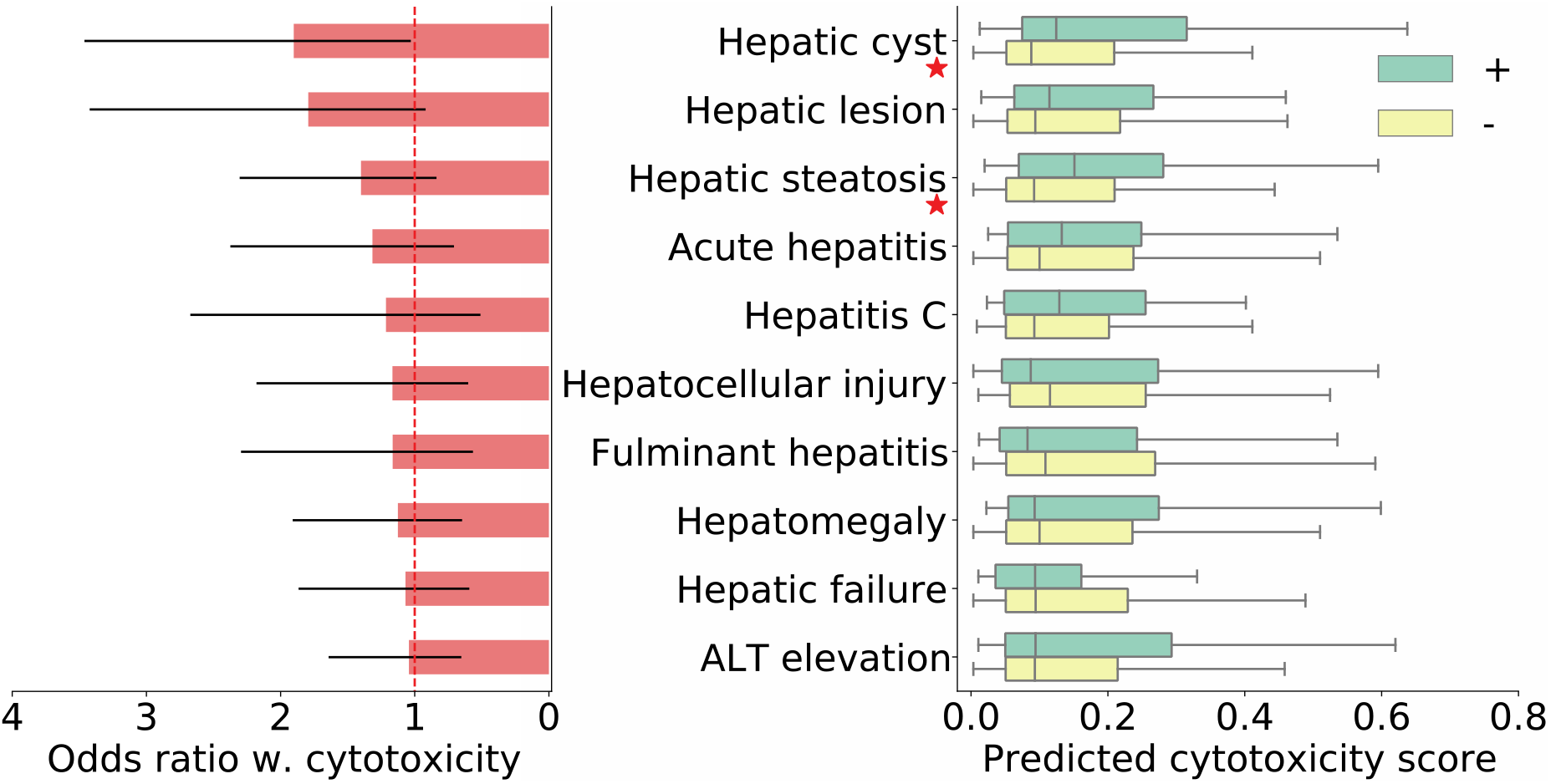
Application of predicted cytotoxicity score in clinical hepatic phenotypes. Boxplot on the right compares the predicted HepG2 cytotoxicity scores among drugs associated with clinical hepatic phenotypes (green box) versus negative controls (yellow box), while barplot on the left shows the odds ratio between HepG2 cytotoxicity and each phenotype (95% confidence interval shown as error bar). Results for ten phenotypes with odds ratio > 1 are shown in the plot. Mann-Whitney U test is employed to examine whether the drugs associated with each phenotype are predicted with higher cytotoxicity scores than the negative controls (red star next to the phenotype name denotes P < 0.05).

Similarly, we sought to differentiate DSSTox compounds associated with drug-induced kidney injury (DIKI) phenotypes from negative controls using predicted HEK293 cytotoxicity score (**Supplementary Fig. 6**). Unfortunately, we were not able to detect significant difference for any of the 24 DIKI phenotypes, as none of them exhibits a significant association with HEK293 cytotoxicity (lower bound of 95% *OR* CI < 1).

## DISCUSSION

Biologically informed VNNs provide a solution to the dilemma posed by conventional supervised learning models: Whether to achieve good predictive performance or high model interpretability. Here, we have explored the implementation of VNNs for predicting and explaining compound response to toxicity assays. Compared to previous efforts, our DTox VNN model uniquely stands out in four aspects: First, the structure of DTox VNN can be customized for an outcome of interest according to the underlying biological processes, making the model flexible towards various assay outcomes. Our hyperparameter tuning process can help with removing unrelated pathways from the network hierarchy and thus adds credibility to model interpretation. Second, trimming of network hierarchy significantly reduces the number of trainable parameters in DTox VNNs, which in turn prevents overfitting. Through comparisons with well-established classification algorithms, we have demonstrated that DTox VNN is a highly efficient learning model with good predictive performance. For instance, DTox VNN achieved the same level of performance as a matched MLP with only three percent of the network parameters. Third, the introduction of an early stopping criterion combined with a relatively small network size makes DTox VNN a fast-learning model. In this study, a learning task that involves 5,000 samples typically converges within 100-200 epochs, with each epoch taking ~90s to complete on a single CPU. Last, and most importantly, DTox advances a novel interpretation framework that identifies high-relevance VNN paths for explaining the toxicity outcome of compounds. The framework builds on top of layer-wise relevance propagation^39^ and assigns a relevance score to each VNN path. The innovation of our framework resides in its ability to statistically assess the significance of each path, with an empirical *p*-value computed from permutation test. Through mechanistic interpretation on individual assays, we have demonstrated the biological significance behind the VNN paths identified by our framework. For instance, in three nuclear receptor assays, our framework was able to consistently identify the corresponding “ground truth” VNN path that represents mechanism of transcription activation. In HepG2 cell viability assay, our framework was able to differentiate distinctive mechanisms leading to cytotoxicity. In aromatase inhibitor and pregnane X receptor agonist assays, our framework was able to disproportionately identify VNN paths that represent the cellular activities induced by specific compounds of interest, implying its potential to detect mechanism of action.

Besides the expected results, DTox also generated new mechanistic hypotheses along model interpretation, some of which can be supported by previous studies. For instance, our framework suggested a potential role for p53 in the regulation of aromatase in MCF-7aro/ERE, a breast cancer cell line. It has been revealed that p53 can directly bind to proximal promoter PII in breast adipose stromal cells, which in turn inhibits aromatase expression^26^. In another case, our framework suggested three signaling pathways, including MAPK/ERK (i.e., MAPK1/ MAPK3 Reactome pathway), Hippo, and TLR3/4, altogether contribute to the HepG2 cytotoxicity of mifepristone. Accordingly, recent studies have pointed out the effect of mifepristone on ERK activation^28^, YAP (a core factor of Hippo) activation^29^, and TLR4 regulation^30^. Particularly, ERK activation by mifepristone can lead to cytotoxicity in uterine natural killer cells^28^ while YAP activation by mifepristone can induce hepatomegaly in mice^29^. Two additional findings from our cytotoxicity analysis have been corroborated by previous studies: (i) The involvement of mitochondria in antigen presentation via ATP synthase and mitochondrial calcium uniporter^35^, and (ii) the disruption of TLR3/4 signaling in DILI^36,37^. In addition, some unexpected findings by DTox are worth further investigation, such as the role of immune response in HepG2 cytotoxicity, the role of interleukin 3/14 in regulation of aromatase, the role of caspase 3/7 in MHC class II antigen presentation, etc.

Despite the highlights mentioned above, DTox bears some limitations in its current form. First, as with all deep learning models, DTox VNN requires a time-consuming hyperparameter tuning process before an optimal model can be reached. And as we observed in the analysis (**Fig. 2a**), an optimal setting may greatly improve the predictive performance of DTox VNN. However, the issue can be resolved with implementation of GPU computing. Second, as we observed in performance assessment, shuffling Reactome hierarchy did not significantly attenuate the predicative performance of DTox VNN, suggesting undocumented interactions between pathways may contribute to toxicity prediction. Therefore, future investigation should be conducted how to train a VNN to recognize these interactions. One possible solution is to incorporate stochastic connections between pathways of distinct branches during training. Finally, despite the incorporation of pathway ontology, DTox VNN did not significantly outperform other well-established classification algorithms, as most differences are within the 95% confidence interval of performance metrics. We noticed the recent development of toxicology-focused graph database such as ComptoxAI, which provides extensive knowledge on relations among chemicals, genes, assays, as well as many other entities^40^. Such database may help us generate more comprehensive feature profile for model training, and thus improves the predicative performance of DTox.

In the future, we expect the application of DTox in two distinct directions. The first direction is concerned with efficacy or toxicity prediction for virtual screening. As what we have accomplished in the screening of ~700,000 DSSTox compounds for cytotoxicity, DTox VNN can quickly go through large-scale chemical libraries and prioritize compounds for further experimental testing. The second direction is concerned with outcome explanation for generating new hypothesis. As we have shown throughout the study, DTox interpretation framework may detect new mechanism of action for compounds, uncover cellular mechanism for outcomes of interest, and identify new therapeutic targets for diseases.

## METHODS

### Processing Tox21 datasets and inferring feature profile for DTox model training

The Tox21 datasets^1^ contain screening results describing the response of *in vitro* toxicity assays to compounds of interest, including approved drugs, experimental drugs, small molecules, and environmental chemicals. We extracted active and inactive compounds from the screening results of each assay, then removed compounds with inconclusive or ambiguous results. We further removed assays with fewer than 5,000 available compounds, focused our analyses on the remaining 15 assays. To quantify structural properties of compounds, we used *rcdk* package to compute 166 binary MACCS fingerprints that cover most of the interesting physicochemical features for drug discovery^41^. We then implemented TargetTox^25^, a feature selection pipeline trained on compound-target interaction data, to infer the target-binding probability of each compound from its MACCS fingerprints. As a result, we derived a feature profile containing 361 target proteins for assay outcome modeling.

### Constructing DTox VNN with Reactome pathway hierarchy

We designed VNN structure based on the Reactome pathway hierarchy that comprises root biological processes, child-parent pathway relations, and protein-pathway annotations (downloaded in Aug 2019)^23^. To trim the scale of neural network and prevent overfitting, we adopted two hyperparameters to filter Reactome pathways: (i) minimal pathway size (values for tuning: 5, 20) and (ii) root biological process (values for tuning: ‘gene expression’, ‘immune system’, ‘metabolism’, ‘signal transduction’, and all possible combinations among the four, 15 values in total). We selected the four processes due to their broad coverage and direct involvement in cellular mechanism of toxicity. Each pathway is coded as a hidden module with fixed number of neurons. For a pathway *p*, the number is defined by 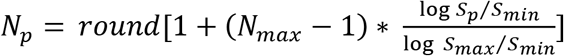, where *S_p_* denotes size of *p*, *S_min_* and *S_max_* denote the minimal and maximal size of pathway in VNN, respectively, *N_max_*(= 20) denotes the maximal number of neurons for a hidden module. As a result, hidden modules of larger pathways are assigned with more neurons to capture potentially more complex response.

Under Reactome hierarchy, DTox VNN model starts from input layer containing 361 protein features, which are connected to lowest-level hidden modules by protein-pathway annotations. The connections to a hidden module of pathway *p* are encoded by a weight matrix ***W_p_*** with dimensions *N_p_*N_protein_*, where *N_p_* denotes the hidden module size, and *N_protein_* denotes the number of input proteins annotated with *p*. With ***W_p_***, input vector ***χ_p_*** is transformed to output vector ***y_p_*** via 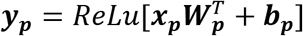, where ***b_p_*** denotes the bias vector. The hidden modules are then interconnected by child-parent pathway relations until root biological processes are reached. Finally, the root biological processes are connected to output layer containing assay outcome. The connections to output layer are encoded by a weight matrix ***W_r_*** with dimensions *1*N_r_*, where *N_r_* denotes the sum of root hidden module sizes. The final output *y_r_* is comupted as 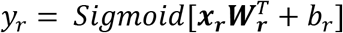. In addition, we adopted the idea of auxiliary layer from DCell^19^ to prevent gradients from vanishing in the lower hierarchy, and to facilitate the learning of new patterns from individual pathways. Specifically, output vector of a hidden module ***y_p_*** is transformed to an auxiliary scalar 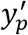 via 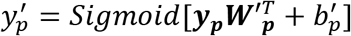, where 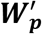 denotes the weights matrix with dimensions 1\**N_p_*. The auxiliary scalars from all hidden modules are then evaluated in a loss function along with the final output: 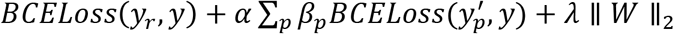. The auxiliary factor α is a hyperparameter of VNN model (values for tuning: 0.1, 0.5, 1), balancing between root and auxiliary loss terms. *β_p_* serves as the adjustment factor for auxiliary loss term from pathway *p*, being computed as the inverse number of pathway count within the corresponding hidden layer. Therefore, pathways in the higher hierarchy exhibit greater contribution to the loss function as pathway count decreases dramatically along the hierarchy. λ (= 1e^-4^) is the coefficient for L2 regularization.

### Learning optimal VNN model for Tox21 assay outcome prediction

Each dataset is split into learning and validation sets by ratio of 4:1. During model training, the learning set is further split into training and testing sets by ratio of 7:1. At every epoch, forward and backward propagation are performed on the training set for deriving gradients of model parameters. The parameters are then optimized by ADAM algorithm with mini-batch size of 32. At the end of every epoch, loss function is evaluated on the testing set for determining the early stopping criterion. Specifically, model training stops if the testing loss has not decreased for 20 epochs.

As mentioned above, DTox VNN model has three hyperparameters: minimal pathway size, root biological process, and the auxiliary factor α. To find the optimal setting for each assay, we adopted grid search and implemented all possible (90 in total; **Supplementary Table 5**) hyperparameter combinations to train VNN models. We evaluated each trained model by computing loss function on the whole learning set, then identified the optimal model that minimizes learning loss. Finally, the held-out validation set was used to evaluate the performance of optimal VNN model and compare with other machine learning models. We adopted two performance metrics for the task: area under the ROC curve (AUROC) and balanced accuracy. We computed the 95% confidence interval (CI) of metrics using bootstrapped samples from predicted outcome probabilities. On average, the bootstrapped samples contain 63.3% of unique original samples. The performance of two methods is significantly different if their CIs do not overlap. Four machine learning models were considered for performance comparison: (i) an alternative VNN model built under shuffled Reactome pathway hierarchy while the shuffle preserves the number of children for each parent pathway and the number of connections between hidden layers (ii) a fully-connected multi-layer perceptron model with the same number of hidden layers and neurons as optimal VNN model, (iii) an optimal random forest model derived from hyperparameter tuning (**Supplementary Table 5**), and (iv) an optimal gradient boosting model derived from hyperparameter tuning (**Supplementary Table 5**).

### Interpretating optimal VNN model by layer-wise relevance propagation

Layer-wise relevance propagation^39^ (LRP) is a model interpretation tool for deep neural networks. Through backward propagation, LRP assigns each neuron a share of the network output, redistributes it to its predecessors in equal amount until input layer is reached. The propagation procedure ensures that relevance conservation is an inherent property of LRP. To implement LRP, we adopted two local propagation rules: generic rule and input-layer rule^42^.

Generic rule was applied to relevance propagation of the hidden neurons. For two connected neurons *j* and *k* from a child-parent pathway pair, the forward propagation of VNN follows *a_k_* = *ReLu*(*∑_j_a_j_w_jk_* + *b_k_*), where *a_k_* denotes the activation of neuron *k*. The generic rule propagates relevance between them as 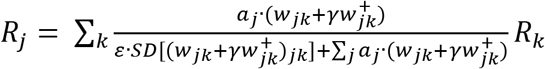, where *γ* and *ε* are two hyperparameters of the rule. *γ* (values for tuning: 0.001, 0.01, 0.1) controls the contribution of positive weights in relevance propagation. Increasing the value of *γ* can marginalize neurons with negative weights and decrease the variance of relevance across neurons, and thus may lead to more stable interpretation results. *ε* (values for tuning: 0.001, 0.01, 0.1) absorbs relevance from neurons with weak or contradictory weights. Increasing the value of *ε* can give prominence to a few neurons with high weights, and thus may lead to more sparse interpretation results.

Input-layer rule was only applied to relevance propagation of the input protein features. For a protein feature *i* and its connected neuron *j* from a lowest-level pathway, the input-layer rule propagates relevance between them as 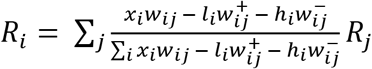, where *l_i_*(= 0) and *h_i_* (= 1) are the lower and upper bound of input feature values.

### Identifying significant VNN paths for explaining toxicity outcome of compounds

After relevance of each neuron being assigned via LRP, a relevance score is computed for each pathway by summing up the relevance scores of its neurons. An observed score is then computed for each VNN path connecting input protein feature to output assay outcome as: 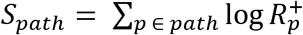, where *p* denotes a protein or pathway along the path. The relevance scores are converted to non-negative values, as we are only interested in the proteins or pathways that are more likely to result in toxicity outcome. The log transformation is adopted to adjust the scale of relevance scores from different layers, as the number of pathways decreases dramatically along the hierarchy.

To assess the significance of each observed path score, we employed a permutation-based strategy to derive the null distribution. Specifically, we shuffled the outcome label of each Tox21 dataset, then re-trained random VNN models using the same hyperparameter setting as previously trained optimal model. The procedure was repeated for *N* = 200 times, a balance between sample size and running time. Scores derived from the random VNN models comprise the null distribution for each observed path score, and thus the empirical P-value can be computed as 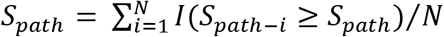. We used false discovery rate (FDR) to perform multiple testing correction on all VNN paths, then identified the significant paths (FDR < 0.05) for each active compound.

As mentioned above, DTox interpretation framework has two hyperparameters: *γ* and *ε* from the generic rule. To study the effect of hyperparameter settings on model interpretation, we implemented all possible (9 in total) hyperparameter combinations to identify significant VNN paths for active compounds. We measured the similarity between each pair of settings by the median Jaccard Index among active compounds regarding their identified significant paths.

### Processing LINCS dataset for validation of DTox interpretation results

The LINCS dataset^24^ contains gene-expression profiles derived after genetic and small-molecule perturbations on a number of cell lines, including MCF-7 (cell line of aromatase assay) and HepG2 (cell line of mitochondria toxicity assay, PXR agonist assay, and HepG2 cell viability assay). We extracted the profiles induced by active compounds of the four assays in their respective cell line. We removed the profiles that did not pass quality control, then separated the remaining ones into three groups based on dose and time of perturbation (1.11μM-24h, 10μM-6h, 10μM-24h). We used the LINCS level 5 data, which consists of moderated differential expression Z-scores, for the validation analysis.

To assess the differential expression of VNN paths identified for each compound, we first identified differentially expressed genes (DEGs) from the corresponding profile by |Z| > 2, as suggested by LINCS. Then, we used Fisher’s exact test to examine whether the pathways along each VNN path are enriched for DEGs. A test *p*-value was computed for each pathway. We used FDR to perform multiple testing correction on all pathways along each path. A VNN path is differentially expressed if all the pathways involved are significantly enriched for DEGs (FDR < 0.05). Finally, we calculated the proportion of differentially expressed paths among the paths identified by DTox (observed proportion) and among all possible paths in VNN (expected proportion).

### Processing NSIDES dataset for analyzing DTox results on HepG2- and HEK293-cytotoxic compounds

The NSIDES dataset^43^ contains drug-adverse event relations that are derived from FDA reports after adjusting for confounding factors. Each drug-adverse event pair is assigned with a proportional reporting ratio (PRR) score along with its 95% CI, which measures the extent to which the adverse event is disproportionately reported among individuals taking the drug. We manually curated a list of 20 clinical phenotype terms associated with drug-induced liver injury (DILI; **Supplementary Table 6**) and a list of 24 clinical phenotype terms associated with drug-induced kidney injury (DIKI; **Supplementary Table 6**). Drugs associated with each phenotype of interest are identified by the lower bound of 95% CI (> 1). Drugs not associated with each phenotype of interest (negative controls) are identified by both the lower (< 1) and the upper (> 1) bound of 95% CI.

To measure the association between each DILI phenotype and HepG2 cytotoxicity, we calculated the odds ratio and its 95% CI based on a 2*2 contingency table. The same procedure was performed to measure the association between each DIKI phenotype and HEK293 cytotoxicity. We also used Fisher’s exact test to evaluate the enrichment of nine cell death-related pathways among the drugs associated with DILI phenotypes. The odds ratio and test P-value were computed for each phenotype-pathway pair. We used FDR to perform multiple testing correction on all phenotype-pathway pairs.

## ACKNOWLEDGEMENTS

This work was supported by NIH grants P30 ES013508, R01 LM010098, R01 AG066833, and K99 LM013646.

## COMPLETING FINANCIAL INTERESTS

The authors declare no competing financial interest.

## AUTHOR CONTRIBUTIONS

J.H.M. and Y.H. conceived the DTox project. J.H.M. and Y.H. designed the DTox model and data analysis workflow. Y.H. and J.D.R performed the analysis. J.H.M. and Y.H. interpreted the results and wrote the paper with editing by J.D.R. All authors read and approved the final manuscript.

## REFERENCES

1 Richard, A. M. et al. The Tox21 10K Compound Library: Collaborative Chemistry Advancing Toxicology. Chem Res Toxicol 34, 189–216, doi:10.1021/acs.chemrestox.0c00264 (2021).

2 Kleinstreuer, N. C. et al. Phenotypic screening of the ToxCast chemical library to classify toxic and therapeutic mechanisms. Nature biotechnology 32, 583–591, doi:10.1038/nbt.2914 (2014).

3 Huang, R. et al. Modelling the Tox21 10 K chemical profiles for in vivo toxicity prediction and mechanism characterization. Nat Commun 7, 10425, doi:10.1038/ncomms10425 (2016).

4 Chan, H. C. S., Shan, H., Dahoun, T., Vogel, H. & Yuan, S. Advancing Drug Discovery via Artificial Intelligence. Trends in pharmacological sciences 40, 592–604, doi:10.1016/j.tips.2019.06.004 (2019).

5 Hemmerich, J. & Ecker, G. F. In silico toxicology: From structure–activity relationships towards deep learning and adverse outcome pathways. Wiley Interdisciplinary Reviews: Computational Molecular Science 10, e1475 (2020).

6 Sedykh, A. et al. Use of in vitro HTS-derived concentration-response data as biological descriptors improves the accuracy of QSAR models of in vivo toxicity. Environ Health Perspect 119, 364–370, doi:10.1289/ehp.1002476 (2011).

7 Liu, J. et al. Predicting hepatotoxicity using ToxCast in vitro bioactivity and chemical structure. Chem Res Toxicol 28, 738–751, doi:10.1021/tx500501h (2015).

8 Ammad-ud-din, M. et al. Integrative and personalized QSAR analysis in cancer by kernelized Bayesian matrix factorization. J Chem Inf Model 54, 2347–2359, doi:10.1021/ci500152b (2014).

9 Yamane, J. et al. Prediction of developmental chemical toxicity based on gene networks of human embryonic stem cells. Nucleic acids research 44, 5515–5528, doi:10.1093/nar/gkw450 (2016).

10 Capuzzi, S. J., Politi, R., Isayev, O., Farag, S. & Tropsha, A. QSAR Modeling of Tox21 Challenge Stress Response and Nuclear Receptor Signaling Toxicity Assays. Frontiers in Environmental Science 4, doi:10.3389/fenvs.2016.00003 (2016).

11 Zhang, J., Mucs, D., Norinder, U. & Svensson, F. LightGBM: An Effective and Scalable Algorithm for Prediction of Chemical Toxicity-Application to the Tox21 and Mutagenicity Data Sets. J Chem Inf Model 59, 4150–4158, doi:10.1021/acs.jcim.9b00633 (2019).

12 Mayr, A., Klambauer, G., Unterthiner, T. & Hochreiter, S. DeepTox: Toxicity Prediction using Deep Learning. Frontiers in Environmental Science 3, doi:10.3389/fenvs.2015.00080 (2016).

13 Idakwo, G. et al. Deep Learning-Based Structure-Activity Relationship Modeling for Multi-Category Toxicity Classification: A Case Study of 10K Tox21 Chemicals With High-Throughput Cell-Based Androgen Receptor Bioassay Data. Frontiers in physiology 10, 1044, doi:10.3389/fphys.2019.01044 (2019).

14 Matsuzaka, Y. & Uesawa, Y. Molecular Image-Based Prediction Models of Nuclear Receptor Agonists and Antagonists Using the DeepSnap-Deep Learning Approach with the Tox21 10K Library. Molecules 25, doi:10.3390/molecules25122764 (2020).

15 Polishchuk, P. Interpretation of Quantitative Structure-Activity Relationship Models: Past, Present, and Future. J Chem Inf Model 57, 2618–2639, doi:10.1021/acs.jcim.7b00274 (2017).

16 Ribeiro, M. T., Singh, S. & Guestrin, C. in Proceedings of the 22nd ACM SIGKDD International Conference on Knowledge Discovery and Data Mining - KDD’16 1135–1144 (2016).

17 Shrikumar, A., Greenside, P. & Kundaje, A. in International conference on machine learning. 3145–3153 (PMLR).

18 Du, M., Liu, N. & Hu, X. Techniques for interpretable machine learning. Communications of the ACM 63, 68–77, doi:10.1145/3359786 (2019).

19 Ma, J. et al. Using deep learning to model the hierarchical structure and function of a cell. Nature methods 15, 290–298, doi:10.1038/nmeth.4627 (2018).

20 Kuenzi, B. M. et al. Predicting Drug Response and Synergy Using a Deep Learning Model of Human Cancer Cells. Cancer cell 38, 672–684 e676, doi:10.1016/j.ccell.2020.09.014 (2020).

21 Lin, C. H. & Lichtarge, O. Using Interpretable Deep Learning to Model Cancer Dependencies. Bioinformatics, doi:10.1093/bioinformatics/btab137 (2021).

22 Elmarakeby, H. A. et al. Biologically informed deep neural network for prostate cancer discovery. Nature 598, 348–352, doi:10.1038/s41586-021-03922-4 (2021).

23 Jassal, B. et al. The reactome pathway knowledgebase. Nucleic acids research 48, D498–D503, doi:10.1093/nar/gkz1031 (2020).

24 Subramanian, A. et al. A Next Generation Connectivity Map: L1000 Platform and the First 1,000,000 Profiles. Cell 171, 1437–1452 e1417, doi:10.1016/j.cell.2017.10.049 (2017).

25 Hao, Y. & Moore, J. H. TargetTox: A Feature Selection Pipeline for Identifying Predictive Targets Associated with Drug Toxicity. J Chem Inf Model 61, 5386–5394, doi:10.1021/acs.jcim.1c00733 (2021).

26 Wang, X. et al. Prostaglandin E2 inhibits p53 in human breast adipose stromal cells: a novel mechanism for the regulation of aromatase in obesity and breast cancer. Cancer Res 75, 645–655, doi:10.1158/0008-5472.CAN-14-2164 (2015).

27 Iorga, A. & Dara, L. Cell death in drug-induced liver injury. Adv Pharmacol 85, 31–74, doi:10.1016/bs.apha.2019.01.006 (2019).

28 Chen, Y., Wang, Y., Zhuang, Y., Zhou, F. & Huang, L. Mifepristone increases the cytotoxicity of uterine natural killer cells by acting as a glucocorticoid antagonist via ERK activation. PloS one 7, e36413, doi:10.1371/journal.pone.0036413 (2012).

29 Yao, X. P. et al. PXR mediates mifepristone-induced hepatomegaly in mice. Acta Pharmacol Sin 43, 146–156, doi:10.1038/s41401-021-00633-4 (2022).

30 Srivastava, M. D., Thomas, A., Srivastava, B. I. & Check, J. H. Expression and modulation of progesterone induced blocking factor (PIBF) and innate immune factors in human leukemia cell lines by progesterone and mifepristone. Leuk Lymphoma 48, 1610–1617, doi:10.1080/10428190701471999 (2007).

31 Brentnall, M., Rodriguez-Menocal, L., De Guevara, R. L., Cepero, E. & Boise, L. H. Caspase-9, caspase-3 and caspase-7 have distinct roles during intrinsic apoptosis. BMC Cell Biol 14, 32, doi:10.1186/1471-2121-14-32 (2013).

32 Yosefzon, Y. et al. Caspase-3 Regulates YAP-Dependent Cell Proliferation and Organ Size. Mol Cell 70, 573–587 e574, doi:10.1016/j.molcel.2018.04.019 (2018).

33 Wang, C. & Youle, R. J. The role of mitochondria in apoptosis*. Annu Rev Genet 43, 95–118, doi:10.1146/annurev-genet-102108-134850 (2009).

34 Albensi, B. C. What Is Nuclear Factor Kappa B (NF-kappaB) Doing in and to the Mitochondrion? Front Cell Dev Biol 7, 154, doi:10.3389/fcell.2019.00154 (2019).

35 Bonifaz, L., Cervantes-Silva, M., Ontiveros-Dotor, E., Lopez-Villegas, E. & Sanchez-Garcia, F. A Role For Mitochondria In Antigen Processing And Presentation. Immunology, doi:10.1111/imm.12392 (2014).

36 Yin, S. & Gao, B. Toll-like receptor 3 in liver diseases. Gastroenterol Res Pract 2010, doi:10.1155/2010/750904 (2010).

37 Guo, J. & Friedman, S. L. Toll-like receptor 4 signaling in liver injury and hepatic fibrogenesis. Fibrogenesis Tissue Repair 3, 21, doi:10.1186/1755-1536-3-21 (2010).

38 Grulke, C. M., Williams, A. J., Thillanadarajah, I. & Richard, A. M. EPA’s DSSTox database: History of development of a curated chemistry resource supporting computational toxicology research. Comput Toxicol 12, doi:10.1016/j.comtox.2019.100096 (2019).

39 Bach, S. et al. On Pixel-Wise Explanations for Non-Linear Classifier Decisions by Layer-Wise Relevance Propagation. PloS one 10, e0130140, doi:10.1371/journal.pone.0130140 (2015).

40 Romano, J. D., Hao, Y. & Moore, J. H. Improving QSAR Modeling for Predictive Toxicology using Publicly Aggregated Semantic Graph Data and Graph Neural Networks. Pac Symp Biocomput 27, 187–198 (2022).

41 Cereto-Massague, A. et al. Molecular fingerprint similarity search in virtual screening. Methods 71, 58–63, doi:10.1016/j.ymeth.2014.08.005 (2015).

42 Montavon, G., Binder, A., Lapuschkin, S., Samek, W. & Müller, K.-R. Layer-wise relevance propagation: an overview. Explainable AI: interpreting, explaining and visualizing deep learning, 193–209 (2019).

43 Tatonetti, N. P., Patrick, P. Y., Daneshjou, R. & Altman, R. B. Data-driven prediction of drug effects and interactions. Science translational medicine 4, 125ra131–125ra131 (2012).

