## Supplementary Fig for "Knowledge-guided deep learning models of drug toxicity improve interpretation"

### SUPPLMENTARY FIGURES

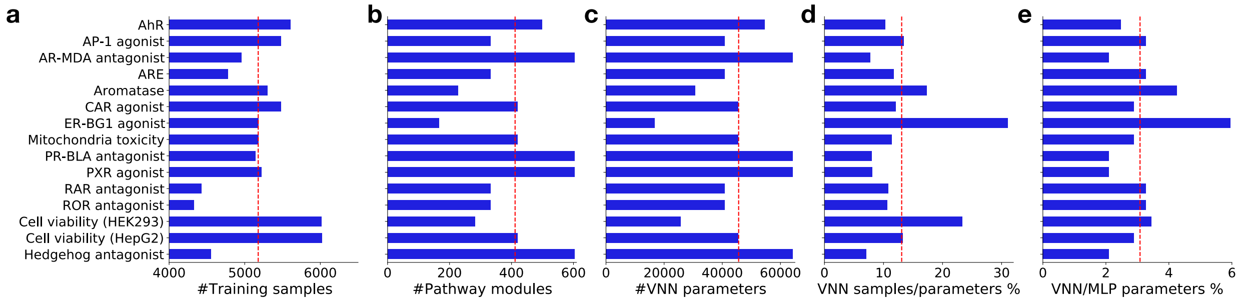


**Suppl. Fig. 1 | Comparison of DTox VNN model statistics**

Barplots showing the comparison of different model statistics for 15 toxicity assays: (**a**) the number of compounds in the training set, (**b**) the number of hidden pathway modules in the optimal DTox VNN model, (**c**) the number of trainable parameters in the optimal DTox VNN model, (**d**) the ratio between number of compounds in the training set versus number of trainable parameters in the optimal DTox VNN model, and (**e**) the ratio between number of trainable parameters in the optimal DTox VNN model versus the matched MLP model. The dashed red line in each panel represents the average across all 15 assays.

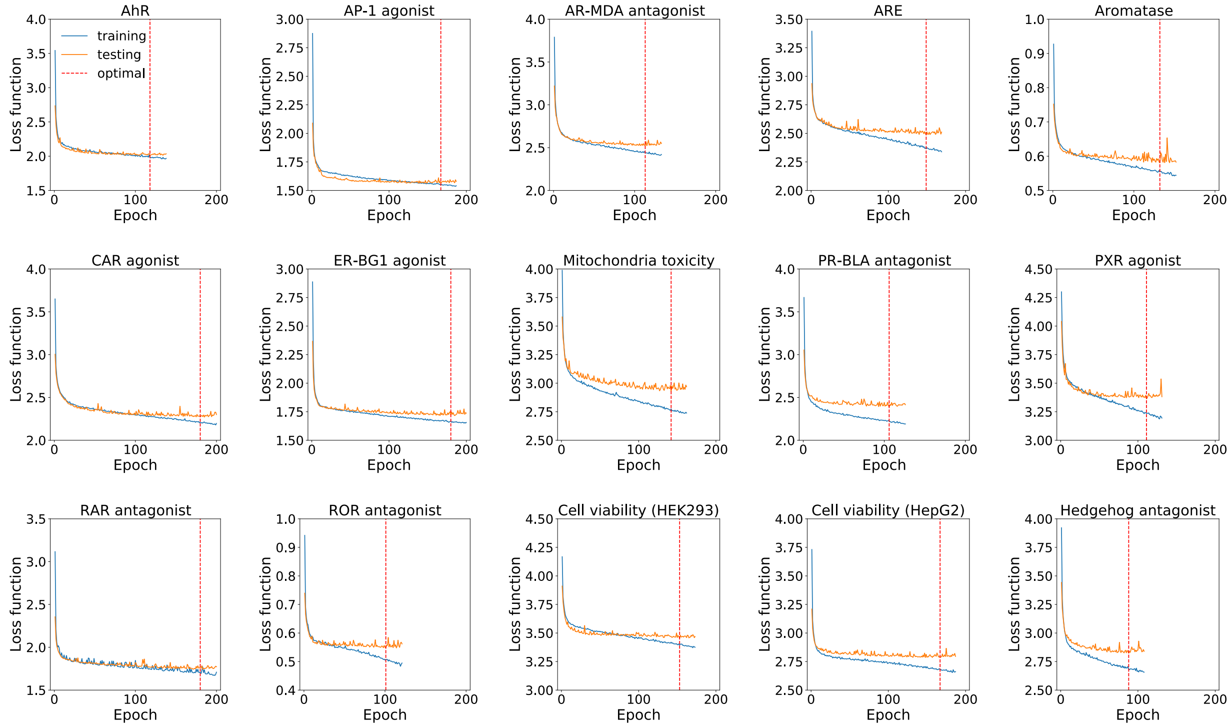


**Suppl. Fig. 2 | Evolution of loss function during learning of optimal DTox VNN models**

Line charts showing the evolution of loss function over epochs during learning process of optimal DTox VNN models for 15 toxicity assays. Two types of loss functions are calculated and shown: loss on the training set (blue line, labeled as training) and loss on the testing set (orange line, labeled as testing). The dashed red line in each chart represents the epoch when optimal model is reached. The testing loss does not decrease for 20 consecutive epochs after the optimal point. AhR: aryl hydrocarbon receptor, AP-1: activator protein-1, AR-MDA: androgen receptor in MDA-kb2 AR-luc cell line, ARE: antioxidant response element, CAR: constitutive androstane receptor, ER-BG1: estrogen receptor in BG1 cell line, PR-BLA: progesterone receptor in PR-UAS-bla HEK293T cell line, PXR: pregnane X receptor, RAR: retinoid acid receptor, ROR: retinoid-related orphan receptor.

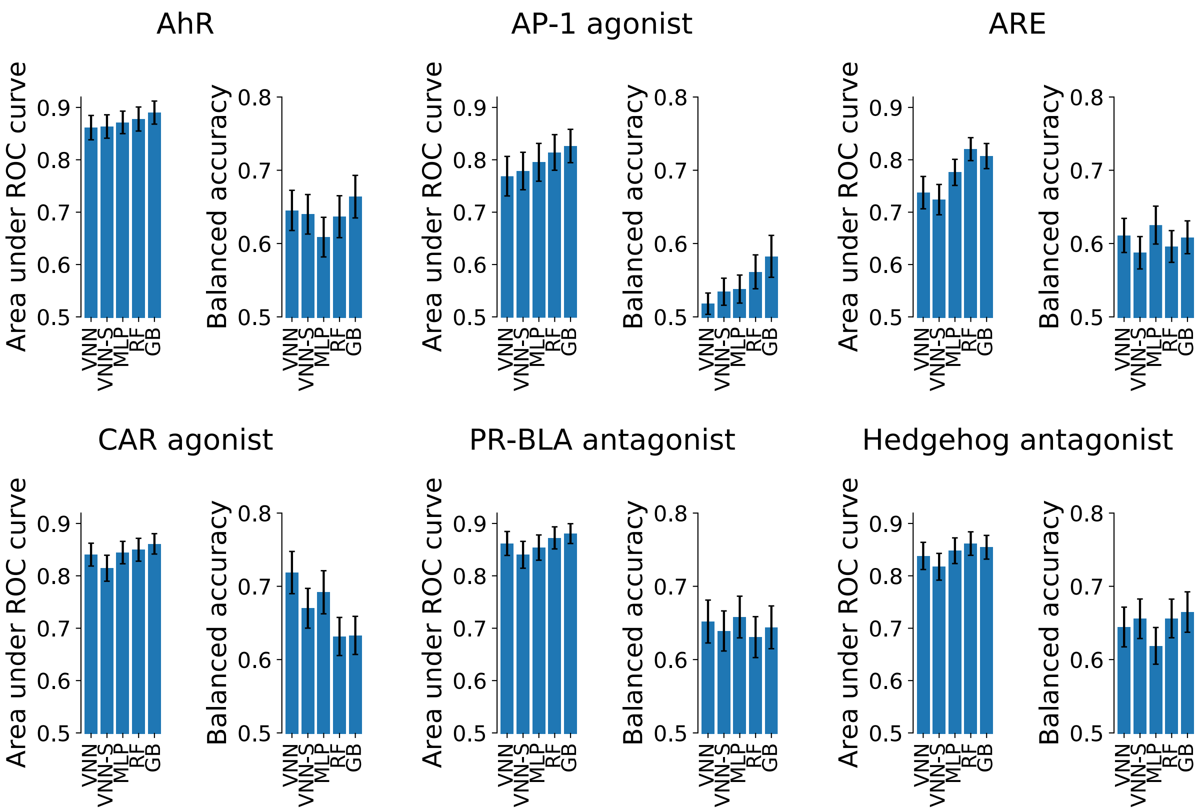


**Suppl. Fig. 3 | Assessment of DTox VNN model performance on held-out validation sets** (related to Fig. 2b)

Barplot showing the validation performance in six toxicity assays (see Fig. 2b for the other nine assays). Performance of DTox VNN (VNN) is compared against four other models: an alternative visible neural network built with shuffled pathway hierarchy (VNN-S), a multi-layer perceptron with the same number of hidden layers and neurons as DTox VNN (MLP), random forest (RF), and gradient boosting (GB). Performance is measured by two metrics: area under ROC curve and balanced accuracy, with error bar shows the 95% confidence interval. AhR: aryl hydrocarbon receptor, AP-1: activator protein-1, ARE: antioxidant response element, CAR: constitutive androstane receptor, PR-BLA: progesterone receptor in PR-UAS-bla HEK293T cell line.

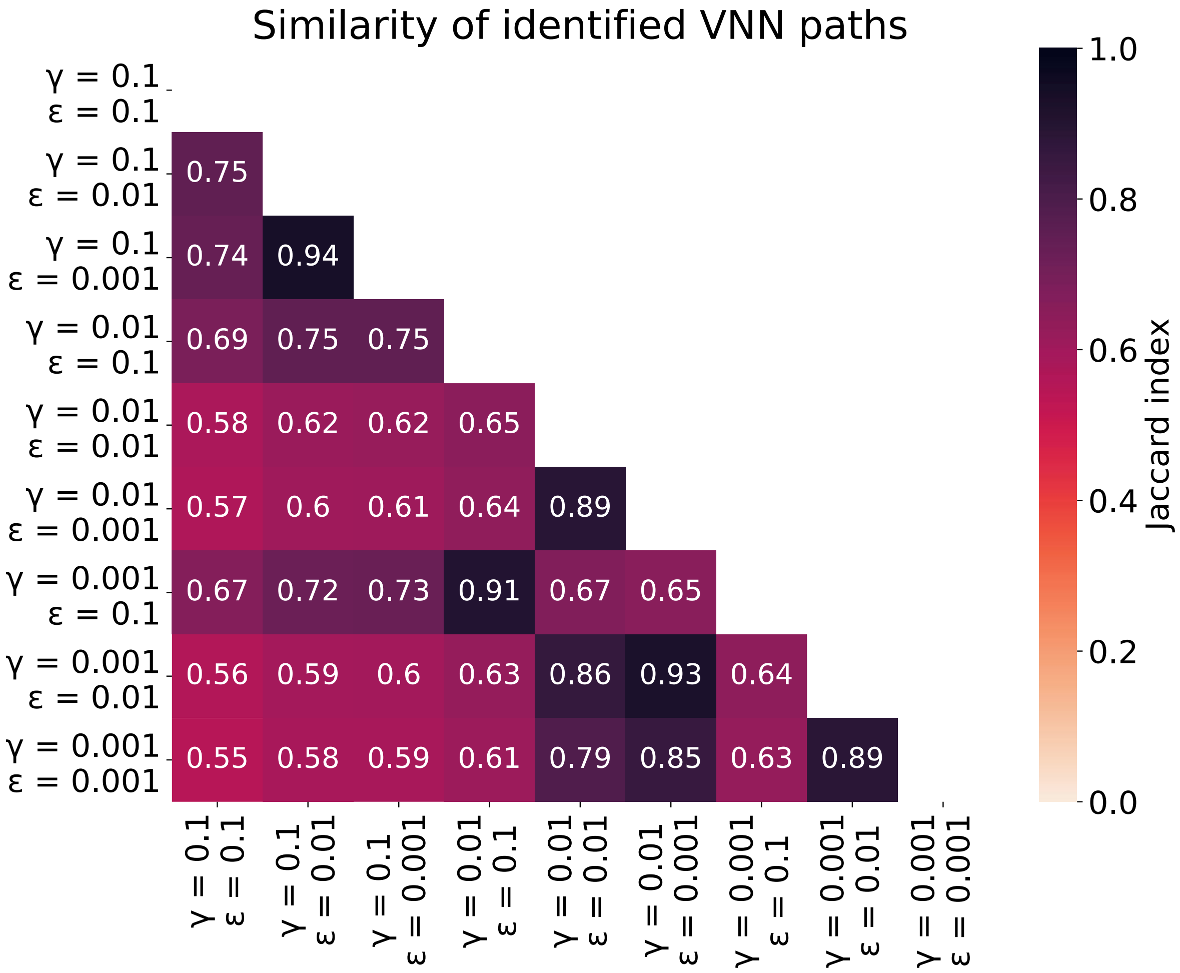


**Suppl. Fig. 4 | Consistency of DTox interpretation across hyperparameter settings**

Heatmap showing the the similarity of VNN paths identified from nine different hyperparameter settings. The similarity between each pair of setting (annotated in each cell) is measured by the median Jaccard Index among active compounds regarding their identified paths.

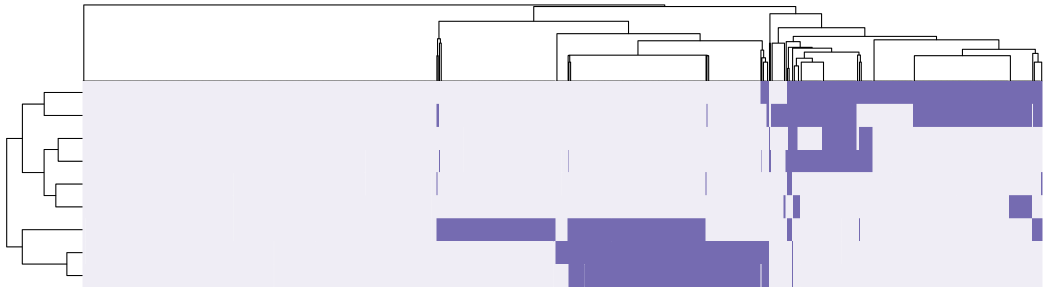


**Suppl. Fig. 5 | Clustering of HepG2-cytotoxic compounds based on cell death-related pathways**

Heatmap showing the mapping between 1,120 HepG2-cytotoxic compounds (columns) and nine cell death-related pathways (rows). Hierarchical clustering is performed for both compounds and pathways. Two clusters appear to form as a result. Compounds in the first cluster (top right) are linked to cytotoxicity via apoptosis-related pathways. Compounds in the second cluster (bottom middle) are linked to cytotoxicity via immune-related and necrosis-related pathway.

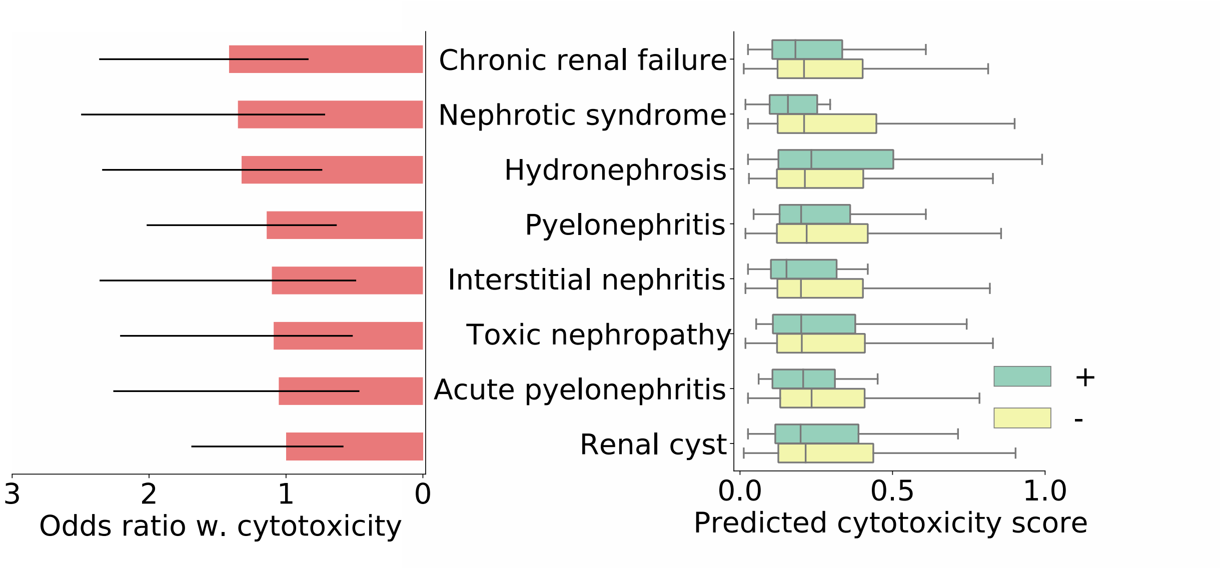


**Suppl. Fig. 6 | Application of predicted cytotoxicity score in clinical renal phenotypes** (related to Fig. 6)

Boxplot on the right compares the predicted HEK293 cytotoxicity scores among drugs associated with clinical renal phenotypes (green box) versus negative controls (yellow box), while barplot on the left shows the odds ratio between HEK293 cytotoxicity and each phenotype (95% confidence interval shown as error bar). Results for eight phenotypes with odds ratio > 1 are shown in the plot. Mann-Whitney U test is employed to examine whether the drugs associated with each phenotype are predicted with higher cytotoxicity scores than the negative controls. No significant signal was detected for any phenotypes.

### SUPPLMENTARY TABLES

In the Excel file, we provide the following six supplementary tables:

**Suppl. Tab. 1: Statistics of DTox VNN models trained on Tox21 datasets**

**Suppl. Tab. 2: Evaluation of model performance on held-out validation Tox21 datasets**

**Suppl. Tab. 3: VNN paths identified for active compounds of Tox21 assays**

**Suppl. Tab. 4: Outcome prediction by DTox for two cell viability assays**

**Suppl. Tab. 5: Hyperparameter tuning of classification algorithms**

**Suppl. Tab. 6: Association statistics between cytotoxicity and liver/kidney injury terms**
